# Reproducible Tract Profiles (RTP2): from diffusion MRI acquisition to clinical practice and research

**DOI:** 10.1101/2022.03.17.484761

**Authors:** Garikoitz Lerma-Usabiaga, Mengxing Liu, Pedro M. Paz-Alonso, Brian A. Wandell

**Affiliations:** Department of Psychology, Stanford University, 450 Serra Mall, Jordan Hall Building, 94305 Stanford, California, USA; BCBL. Basque Center on Cognition, Brain and Language. Mikeletegi Pasealekua 69, Donostia - San Sebastián, 20009 Gipuzkoa, Spain; Wu Tsai Neurosciences Institute, Stanford University, 94305 Stanford, California, USA; IKERBASQUE. Basque foundation for science. 48013 Bilbao, Spain

**Keywords:** Diffusion MRI, White matter tracts, Tractography, Computational reproducibility, Clinical practice

## Abstract

Diffusion MRI is a complex technique, where new discoveries and implementations occur at a fast pace. The expertise needed for data analyses and accurate and reproducible results is increasingly demanding and requires multidisciplinary collaborations. In the present work we introduce Reproducible Tract Profiles (RTP2): a set of flexible and automated methods to analyze anatomical MRI and diffusion weighted imaging (DWI) data for reproducible tractography. The tools read structural MRI data and process them through a succession of serialized containerized analyses. We describe the DWI algorithms used to identify white-matter tracts and their summary metrics, the flexible architecture of the platform, and the tools to programmatically access and control the computations. The combination of these three components provides an easy-to-use automatized tool developed and tested over 20 years, to obtain usable and reliable state-of-the-art diffusion metrics at the individual and group levels for basic research and clinical practice.

**Highlights:**

- Automated, flexible and reproducible protocol for white-matter tractography and tractometry.
- High computational (same data, different computations) and test-retest reproducibility (data from different sessions, different computations).
- Open-source code and publicly available containers.

## Introduction

The complexity of neuroimaging data and analyses challenges researchers’ ability to implement reproducible research methods. In many neuroimaging publications, there is no chance that a reader can replicate the experimental data acquisition or reproduce the computational analyses (Buckheit and Donoho, 1995; Sandve et al., 2013; Wilson et al., 2017). Current practices increasingly invite data sharing among researchers and labs to enable colleagues in the scientific community and clinical practitioners to reproduce, check, and further explore datasets and computational analyses (Peng, 2011; Stodden et al., 2014). Together with the desirable increase of good practices and data sharing there has been a substantial increase in algorithm complexity. Some of this complexity takes the form of a large number of pipeline parameters that users can set. Choices made in these parameters can have substantial effects on the reported MRI results (Botvinik-Nezer et al., 2020; Schilling et al., 2021). Further, it has been pointed out across different MRI techniques that we are often uncertain about critical parameters in computational models (Yarkoni and Westfall, 2017), and DWI and tractography is not an exception. Neuroscientists find it difficult to keep track of the full range of parameters used in any particular analysis, and even fewer neuroscientists record the combinations of parameters they used during data exploration and exploitation (Baker, 2016). These difficulties can be even more common in clinical settings, where clinicians may have access to a large cohort of patients and less time to develop computational research protocols.

There are different available solutions for identifying white matter tracts. Those solutions vary greatly in terms of scope, technical approach, and flexibility. The main difference is based on the level at which they operate. For example, some of them are complete ecosystems where most of the processing steps (preprocessing, registering, modeling and tracking) are included, such as MRtrix (Tournier et al., 2019), FSL (Smith et al., 2004), Tracula (Yendiki et al., 2011), TrackVis (Wang et al., 2007), DSI-studio (Yeh et al., 2010) or Dipy (Garyfallidis et al., 2014). Using any of these solutions, the researcher would be able to obtain their tractography results, and build a pipeline on top of it to automatize most of the tasks. On the other hand, there are some metasolutions that build on top of existing neuroimaging tools, relying on the best available option for image registration, preprocessing, anatomical analyses, diffusion modeling, tractometry, etcetera. In this category, we include TractoFlow (Theaud et al., 2020), TractSeg (Wasserthal et al., 2018) and both the Matlab and Python versions of AFQ (Kruper et al., 2021; Yeatman et al., 2012). Reproducible Tract Profiles (RTP2) falls in this second category.

RTP2 is an automated tool that builds on the state-of-the-art computations of DWI data and tractography and also improves computational reproducibility. The first version of RTP (Lerma-Usabiaga et al., 2019, 2020) was based on a containerization of the matlab-based tool AFQ (Yeatman et al., 2012). RTP2 uses some of the code from the same base repository (https://www.github.com/vistalab/vistasoft), but the tractography system has been redesigned for flexibility and completeness based on Freesurfer (Fischl, 2012), MRtrix (Tournier et al., 2019) and other well-established software tools (see Methods section for details). These methods implement fully reproducible and complete tractography analyses that begin with data acquisition and end with data ready to be input in statistical software. This solution is reproducible because the analysis software and its dependencies are embedded in a container, and the input data, complete set of analysis, configuration parameters and outputs are stored in a single informatics platform. This platform can be a single dedicated computer or a neuroinformatics platform that runs locally, in a server or in the cloud. Together, RTP2 tools comprise a system that enables users to check and reproduce all the computations.

RTP2 has been designed with three objectives: To be (1) reproducible, (2) automated, and (3) flexible. Reproducibility is guaranteed by basing the software on containers that document all of the run-time parameters —see cyan boxes in Figure 1. The system is automated and easy to use, no need for further programming for obtaining the desired tracts. Flexibility is provided by configuration files that allow for any number of ROI and parameter combinations; we provide a default configuration file that enables the user to identify many of the most common tracts out-of-the-box.

**Figure 1.**
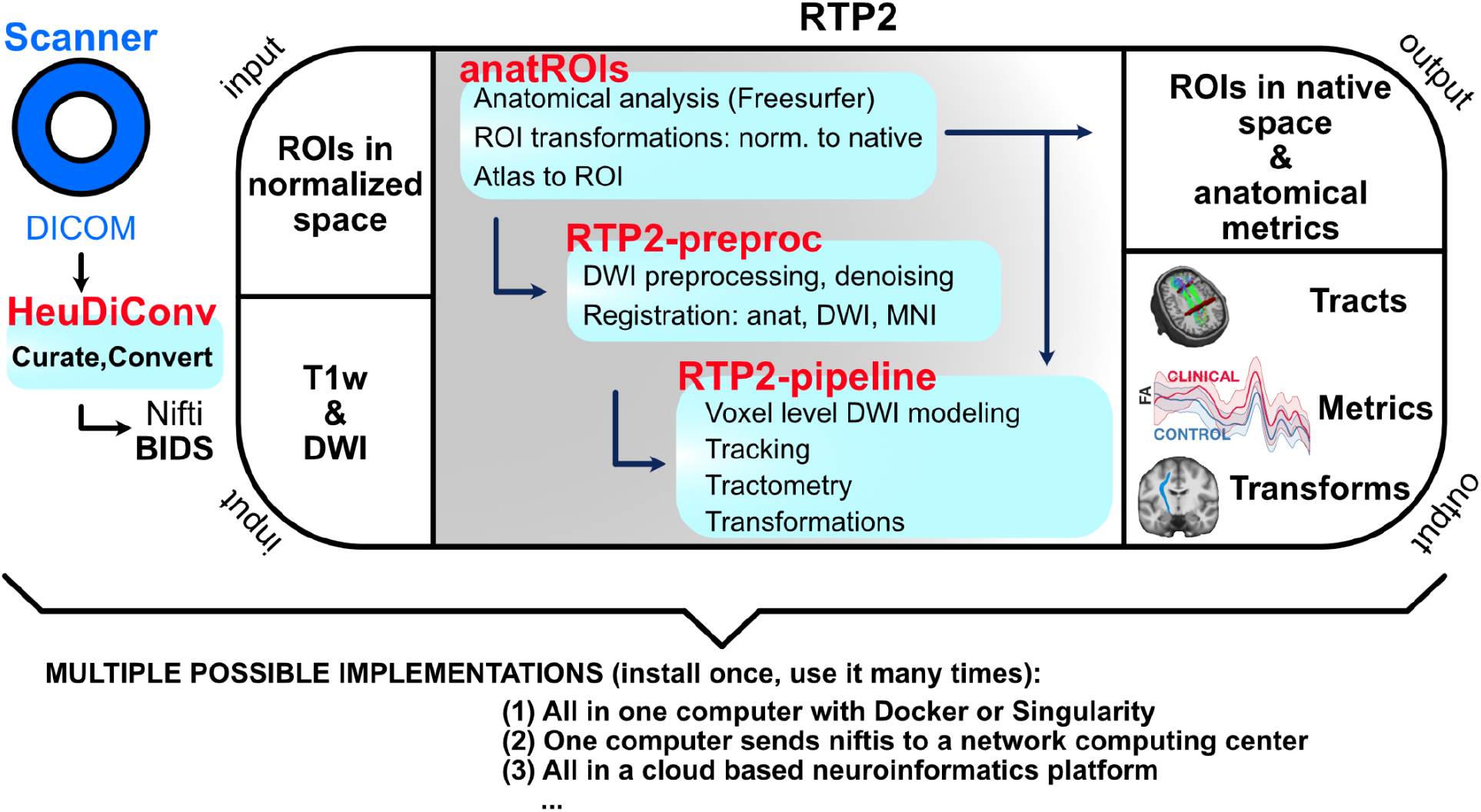
Automated and reproducible pipeline for tractography. RTP2 can be automated to produce reproducible results with minimal human interaction. The DICOMs acquired from the MRI scanner are automatically converted into BIDS format Nifti-s and transferred to the informatics platform. Containerized images containing the anatomical and DWI analysis tools along with files specifying the computational parameters are stored and managed in the system. The system takes individual subject space anatomical images, DWI data and ROIs in normalized space that researchers would like to use in the next steps and coregisters them first. Then it runs the anatomical pipeline and creates structural segmentations and ROIs with Freesurfer, as well as multiple anatomical metrics. The system converts multiple public atlases as well as MNI/surface defined ROIs into volumetric ROIs to be used later in tractography. After the DWI preprocessing, the user can select any combinations of ROIs to define white matter tracts. Tracts can be defined using seed and target ROIs, or can be selected from all the fibers of a whole-brain tractogram (WBT). In the output of the pipeline the user will obtain all the requested tracts and metrics, including along-the-tract profiles (see Figure 5). All structural metrics from Freesurfer as well as all the ROIs in individual-subject space generated from the atlases or input by the researcher will be available. Those ROIs can be used for other analyses, such as fMRI. The whole system can be installed once in a single computer, it can be installed in a distributed system working with high performance computing clusters, or it can be managed in a cloud based neuroinformatic platform.

The system ensures *reproducibility* by basing the computations on containers that include the software and all of its dependencies, and that define the parameters using configuration files that are stored as part of the results. The basic steps are laid out in Figure 1. A container (HeuDiConv) reads the DICOMs from the scanner, converts them to Nifti and curates and transforms them in BIDS format (Gorgolewski et al., 2016). Next, the anatROIs container processes the T1-weighted images using Freesurfer and represents the data in MNI space. The researcher can specify regions-of-interest (ROIs) in normalized space that will be transformed into the individual-subject’s space and that can be used in the next steps of the pipeline. These ROIs can either be selected from several hundred ROIs based on different atlases that are included within RTP2, or they can be created by the researcher in the individual subject’s native space using Nifti format. The RTP2-preproc container processes the diffusion images and registers them with the anatomical images. The RTP2-pipeline container combines the anatomical images, the preprocessed diffusion images and the ROIs to obtain white matter tracts of interest. This container outputs data in a number of ways: tracts in Nifti and tck formats, tracts in volumetric Nifti-s in individual and template space, tract metrics in csv format, and more. The four containers can be installed in a single computer, in a High Performance Cluster (HPC) or in a neuroinformatics platform.

RTP2 is highly *automated* because it is designed for cognitive neuroscientists and/or clinicians interested in conducting research on white matter and tractography. The system quantifies the properties of white-matter tracts at the individual-subject level, and it offers the user high flexibility in terms of how to define white-matter tracts based on white-matter, subcortical and cortical ROIs, as well as the most representative tractrometrics (e.g., FA, RD, AD) and tract-similarity metrics (e.g., Dice,.…). The goal is to help end-users, offering them multiple options that they can choose from. The architecture of the solution, the defaults and the algorithms are explained in the Methods section.

*Flexibility* is provided in two ways: (i) the system can be run in a variety of platforms, depending on the resources available in the lab/clinic; the results will be the same across platforms, and (ii) the user can decide the diffusion model to use, the tracking algorithm, and what to track—with the existing and user created ROIs, the possibilities are practically infinite. In Figure 2 we illustrate all the possible different ROIs RTP2 takes to define tracts. The ROIs can be classical anatomically defined white-matter tracts (Figure 2A), which can be used as targets and seeds themselves, or can be used to select fibers from a whole-brain tractogram. The ROIs can be cortical (Figure 2B) or subcortical structures (Figure 2C) as well. All types of ROIs can be combined to obtain any number of tracts. In the first row of Figure 2 we list most of the ROIs already included in RTP2, based on different atlases and tools. Importantly, the system is prepared to input user defined ROIs without any coding, in the second row of Figure 2, we include examples of ROIs that can be created by the user. In the third row we include some example tracts. In Figure 2A we illustrate the white matter ROIs tracking the arcuate fasciculus: first we obtained the whole-brain tractogram, and then used two white matter ROIs to just select those fibers crossing both ROIs. In Figure 2B we track the arcuate fasciculus using cortical ROIs only, using both ROIs bidirectionally as targets and seeds. Next, to illustrate how we can combine subcortical and cortical ROIs, between Figure 2B and Figure 2C we show the optic radiation, tracked using the same bidirectional target-seed method. At last, we use the same method with two subcortical ROIs (Figure 2C) to obtain the dentatothalamic tract.

**Figure 2.**
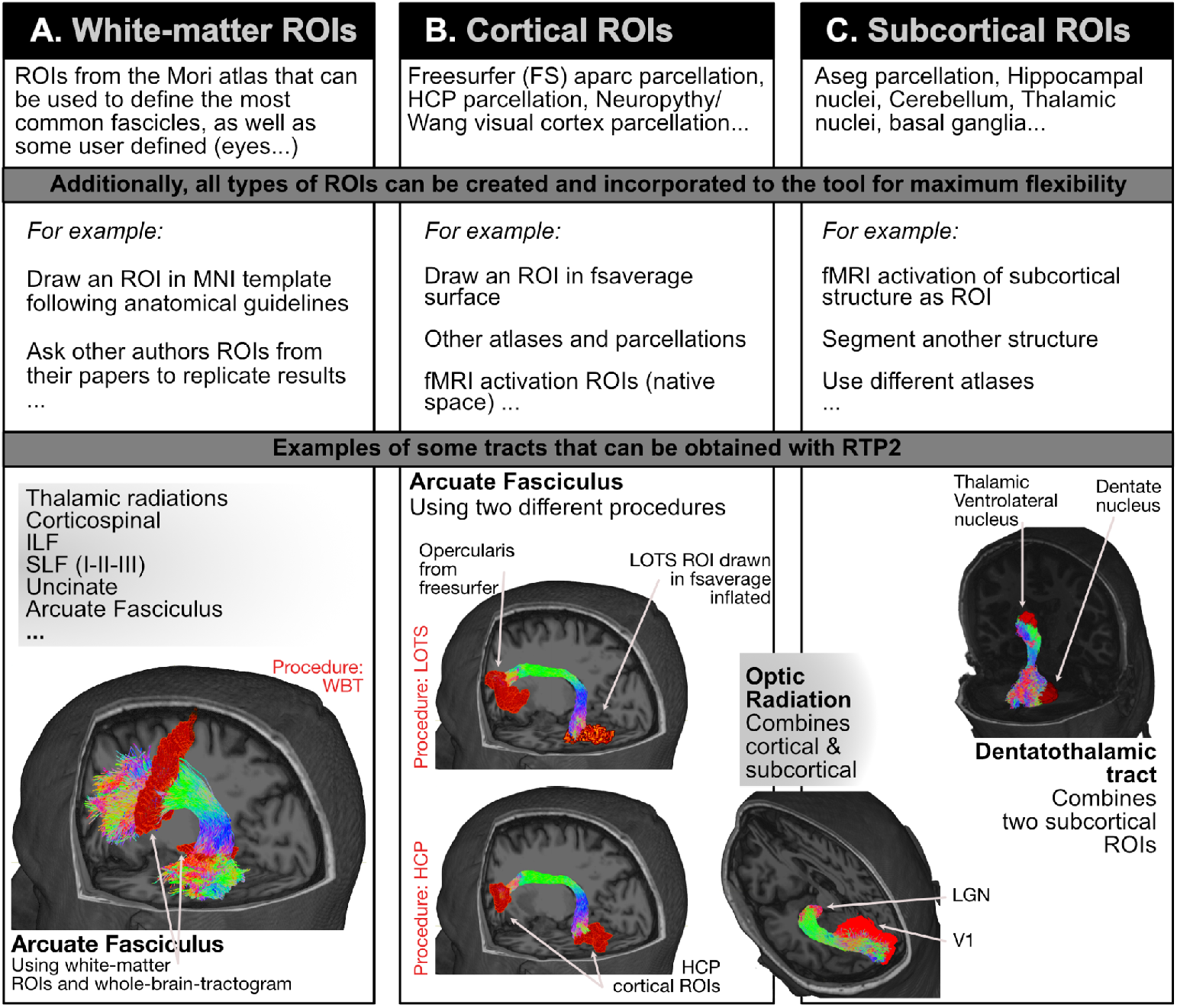
RTP2 Flexibility: any combination of ROIs and several diffusion models can be used to measure tracts of interest. The RTP2 tool allows for an almost infinite combination of ROIs to obtain tracts. Those ROIs can be classical anatomically defined white-matter tracts (A). ROIs are used to select fibers from a whole-brain tractogram. The ROIs can be cortical (B) or subcortical structures (C) as well, and they can be used to track in any combination. Additionally, the system is prepared to input user defined ROIs without any coding (see examples in second row). We include some example tracts that can be obtained from the existing ROIs to illustrate how RTP2 tracking works, but the system is fully configurable by the researcher. In red (WBT, HCP, LOTS) examples for the three procedures used to illustrate the reproducibility analysis.

To characterize the flexibility (and reproducibility) of RTP2, the analytical approach used in the Results section presents a comparison of the computational and test-retest reproducibility of the three different procedures to track the arcuate fasciculus included in Figure 2. The first procedure is based on a 8 million fiber whole-brain tractography (i.e., WBT; marked in red in Figure 2A): we select the fibers that cross two white-matter ROIs to define the arcuate fasciculus. In the second procedure, we use the Human Connectome Project (HCP; in red in Figure 2B) cortical atlas (Van Essen, 2011) to select one ROI corresponding the opercularis region in the inferior-frontal gyrus (IFG) and a second ROI that corresponds to the occipito-temporal sulcus (OTS). We use both ROIs as seeds and targets for tracking the arcuate fasciculus. The third procedure tracks the arcuate fasciculus as well, but we obtain the ROIs differently: the opercularis region is obtained from Freesurfer’s aparc2009 cortical parcellation. The ventral occipito-temporal region ROI is the lateral OTS (LOTS) manually drawn on the inflated surface of Freesurfer’s fsaverage (procedure referred as LOTS in red in Figure 2B). The manually drawn surface ROI is automatically converted to the individual-subject volumetric ROI, in a step included in RTP2. We obtained the profile of FA values along the tract (see Figure 5), as well as the mean value of the whole tract, and compared the values of the three differently tracked arcuate fasciculus. Researchers have different preferences on how to track different tracts, and this comparison illustrates some possibilities in this regard and the differences they might obtain with their data. We obtain very high computational and test-retest reproducibility in most cases. This example illustrates how the researcher can flexibly control and compare between the multiple solutions that can be followed in this tool and of her/his research questions and objectives when tracking which, logically, will impact the final results obtained.

## Results

To illustrate the flexibility and reproducibility of RTP2, we analyzed the bilateral arcuate fasciculus (AF) from a dataset of 112 participants (24 repeated for test-retest) using the three above-mentioned procedures: (1) **WBT**: using a whole-brain-tractogram and white-matter ROIs —Figure 2A, (2) **HCP**: using cortical ROIs obtained from the HCP atlas —Figure 2C,, and (3) **LOTS**: using cortical ROIs obtained from Freesurfer’s atlas and hand-drawn in Freesurfer’s average surface —Figure 2C. We used a double analytical approach: *computational reproducibility*, repeating the computation on the same diffusion data and quantifying changes from computation to computation; and *test-retest reproducibility*, obtaining DWI data from the same subjects and using the same MRI protocol in two different moments. As an initial illustration of the analysis, in Table 1 we show the correlations of the tract’s mean fractional anisotropy (FA; (Basser and Pierpaoli, 1996) values for computational and test-retest measures.

**Table 1.** Evaluation of computational and test-retest reproducibility for arcuate fasciculus mean FA values across three methods.

|  | Computational<br>(r, N=112) | Test-Retest<br>(r, N=24) |
| --- | --- | --- |
| Left WBT (L_WBT) | 0.990 | 0.962 |
| Right WBT (R_WBT) | 0.994 | 0.962 |
| Left HCP (L_HCP) | 0.995 | 0.953 |
| Right HCP (R_HCP) | 0.989 | 0.942 |
| Left LOTS (L_LOTS) | 0.988 | 0.955 |
| Right LOTS (R_LOTS) | 0.984 | 0.933 |

**Figure 3.**
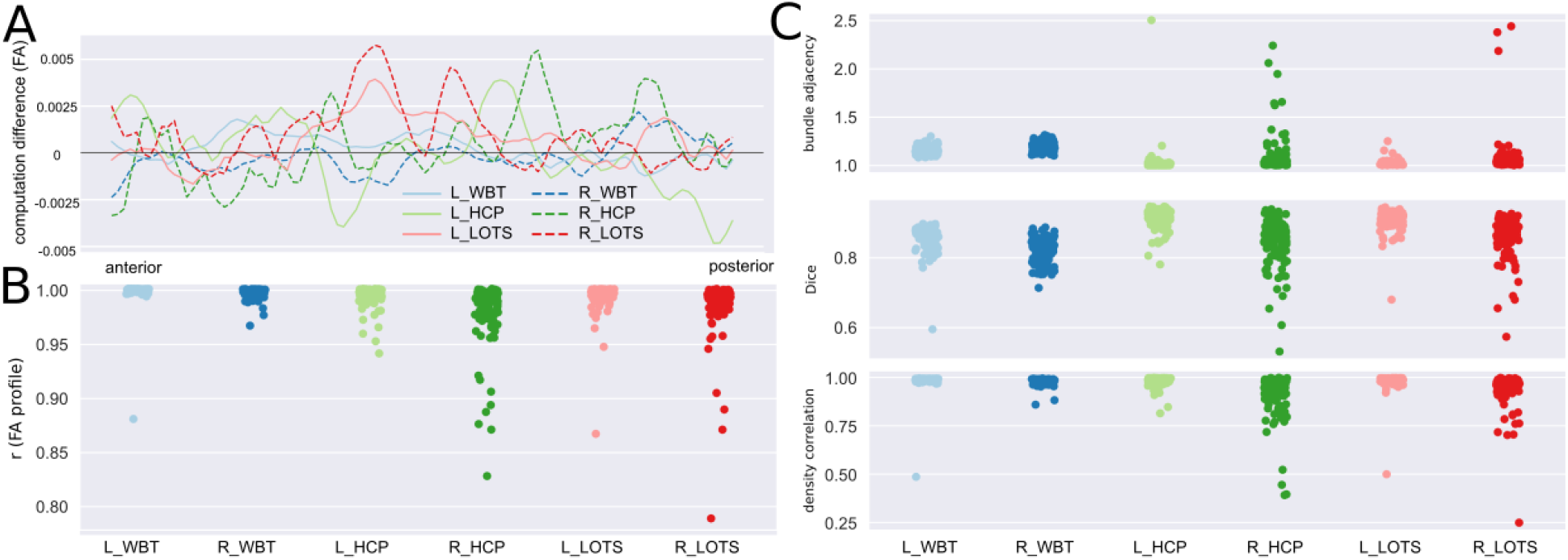
Evaluation of computational reproducibility for arcuate fasciculus tracking across three procedures. **A)** Per procedure/hemisphere across-subject mean FA tract-profile differences between the first and the second computations. **B)** Strip plots showing the distribution of correlation coefficients between all possible pairs computed for each tract and each subject. Each dot represents the correlation coefficient for a specific computation pair for one participant. **C)** Macrostructural agreement indices distribution: bundle adjacency (top), Dice coefficient (middle), and density correlation (bottom) between the two separate computations for each tract and each subject.

Next, we provide computational and test-retest reproducibility analyses based on the whole tract. We show the FA values in the text and other measures in Supplementary Figures.

### Computational reproducibility

To test the computational reproducibility, we repeated the tractography reconstruction using the RTP2-pipeline twice using identical inputs and parameters. For each tract of interest, computational reproducibility was measured with both microstructual (e.g., FA, MD, RD) and macrostructural metrics (e.g., Dice, density correlation, bundle adjacency; Figure 3). The generation of the tract profiles for each tract in RTP2 allows users to conduct further quantitative analysis, not restricted to the mean values of the tract. The computational reproducibility of each individual data point along the tract is good, with FA across-subject average difference between computations being smaller than 0.005 (see Figure 3A). The correlation between the tract profiles for computation 1 and computation 2 tends to be high as well, with the average value in the range of 0.978 ± 0.03 for R_HCP to 0.997 ± 0.01 for R_WBT (Figure 3B).

There are some subjects with relatively lower correlation values in R_HCP and R_LOTS procedures, with correlations ranging from 0.828 to 0.999 and 0.789 to 0.999 respectively at the subject level, and a median across subjects of 0.987 and 0.992. At the macrostructural level, we used several measures to describe reproducibility of streamlines (Figure 3C). This includes (1) volume Dice overlap, which describes the overlap of two tracts, (2) density correlation which describes the similarity of the streamline density of two tracts, (3) bundle adjacency which describes the average distance of disagreement of two tracts. The average Dice overlap ranges from 0.82 ± 0.04 to 0.91 ± 0.03 across the tracts of interest, which reflects that all these tracts have high volume overlap between two separate computations. This high computational reproducibility observed for Dice overlap is also found for the other macrostructural indexes (i.e., density correlation and bundle adjacency) across all the tracts. Similar to the FA value correlations, the R_HCP and R_LOTS procedures showed high variance for the macrostructural measurements, with density correlation values ranging from 0.39 to 0.99 and from 0.25 to 0.99.

### Test-retest reproducibility

The test-retest reproducibility was assessed on a subset of 24 participants, who returned for a second acquisition session within a mean temporal interval of 15 days. The same analytical approaches were adopted to measure test-retest reproducibility using microstructural and macrostructural measurements. Overall, we found high test-retest reproducibility across all the left and right hemisphere white-matter, subcortical and cortical tracts of interest, although some specific tracts showed numerically higher variability. As expected, the reproducibility values are lower in test-retest than in computational reproducibility, with overall higher mean difference between tract profiles in test-retest reproducibility (see Figure 4A). This affects the FA correlation analysis, as well as the coefficients averaged within each tract that ranged from 0.918 ± 0.11 for R_HCP to 0.990 ± 0.01 for L_WBT (see Figure 4B). Again, R_HCP and R_LOTS showed higher numerical variability between test and retest computations, with the subject level FA correlation values for R_HCP ranging from 0.46 to 0.99, and from 0.76 to 0.99 for R_LOTS. At the macrostructural level, the average Dice overlap has a range of 0.78 ± 0.04 to 0.89 ± 0.03 across all the tracts of interest. Consistent with the Dice overlap results, all tracts have high agreement in streamline density and small bundle adjacency. Similar results were observed in terms of variance, with R_HCP and R_LOTS showing numerically higher variance in all indices.

**Figure 4.**
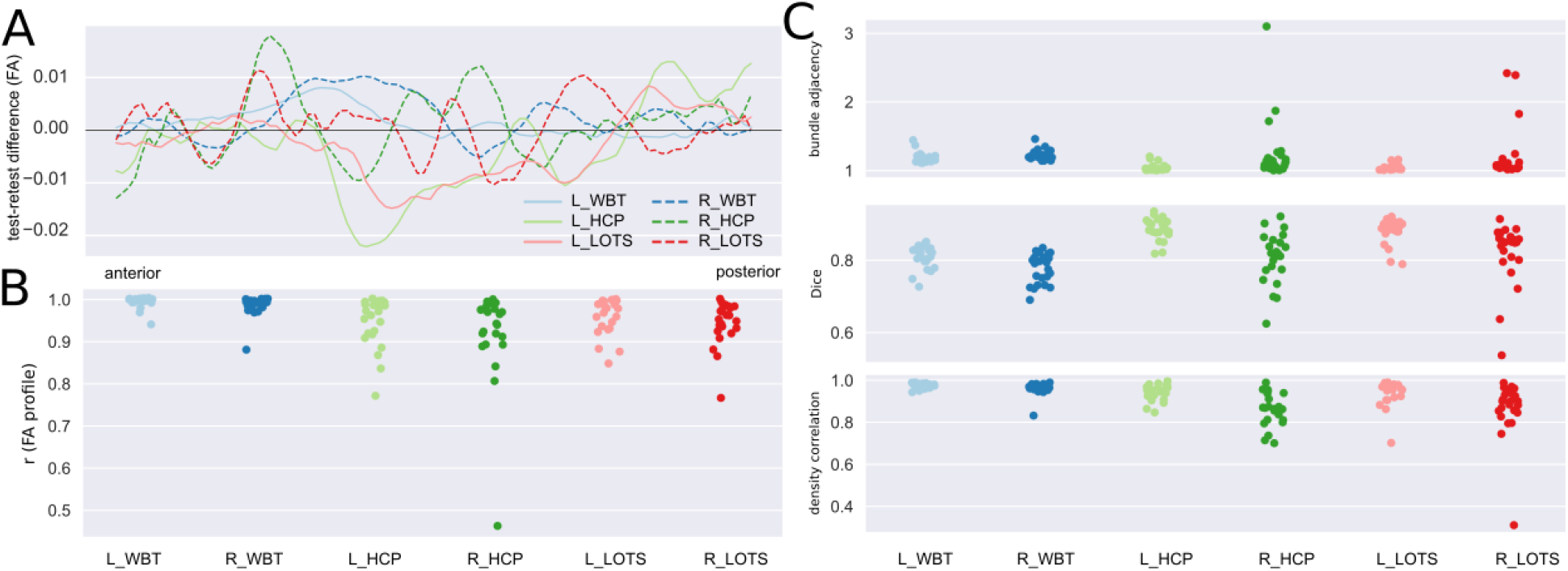
Evaluation of test-retest reproducibility for arcuate fasciculus tracking across three procedures. **A)** Per procedure/hemisphere across-subject mean FA tract-profile differences between the test and retest acquisitions. **B)** Strip plots showing the distribution of the correlation coefficients between test and retest computations for each tract and each subject. **C)** Macrostructural agreement indices distribution: bundle adjacency (top), Dice coefficient (middle), and density correlation (bottom) for test and retest computations for each tract and each subject.

## Discussion

The increasing complexity in diffusion neuroimaging data and analyses challenges researchers’ ability to implement reproducible research methods. The technical and conceptual expertise required to obtain reproducible results is being addressed in the neuroimaging community with the creation of automated tools that enable non-experts to conduct analyses using state-of-the-art methods with better validity and reliability. Additionally, some of the complexity takes the form of a large number of parameters that users can set. Choices made in these parameters can have substantial effects on the reported results, and it is critical to have a system that keeps track of the parameters used in any particular analysis, and to record the combinations of parameters used during data exploration (Baker, 2016). With these objectives in mind we developed a reproducible, automated and flexible system that, once installed, can automatically run the tractography pipeline, and keep track of all the analytical steps performed.

We illustrated the functionality and the reproducibility of the pipeline examining computational reproducibility on 112 subjects and test-retest reproducibility on 24 of those subjects with repeated measures. We obtained nearly perfect computational reproducibility values in all metrics, at the mean tract level as well as at the whole-tract level. We illustrated the flexibility of the tool by computing the same tract, the AF, in three different ways but using the same tool. This is something common in the field: the same anatomical structure can be defined in several different ways (Schilling et al., 2021); and, typically, that occurs using different pipelines and softwares in different laboratories. In our examples, all ROIs came from an average space and were converted to individual-subject space, but RTP2 also allows defining ROIs directly in native space as well, e.g.: as a result from individual subject fMRI experiments. Results revealed that although the computational reproducibility is nearly perfect, it is not absolutely perfect. This is expected, as the whole pipeline contains several non-deterministic calculations that can slightly influence the final results. A fully reproducible pipeline can be achieved by having the random seeding initialization fixed and the interested researcher will be able to choose such models in RTP2. In our analyses we used a probabilistic algorithm to generate the streamlines, as it generally provides the best results for most tracts (Bonilha et al., 2015; Grisot et al., 2021; Khalsa et al., 2014). Additionally, having the random seeding fixed is not compatible with multi-threaded steps. Multi-threading allows for faster computations, but introduces randomness in the order of execution and generally slightly impacts the reproducibility of results as well.

As indicated, RTP2 can be also used in clinical research with clinical populations. For example this tool can be used for surgery preparation (e.g., tracking the dentatothalamic tract in patients with Parkinson disease), and it can be used to further understand the involvement of white-matter tracts in the spreading of grand mal (e.g., patients with intractable epilepsy). RTP2 can be also used in infants with brain tumors, overcoming a triple challenge: (1) low myelination of infant brains, and therefore, weaker diffusion signals, (2) different head sizes, requiring the use of non-standard templates, and (3) non-standard tracts, as the tumors deviate standard natural pathways. Thus, RTP2 is robust enough to work with different clinical populations across the whole life span, and it is flexible enough to allow exploring many different possibilities even in patients with structural alterations. Since RTP2 has no priors embedded, it opens the door to model any shaped tracts at the individual subject level.

This project is open-source, it is available in Github and Dockerhub to any researcher interested in using it. This means as well that anyone can extend the functionalities, making them available to every user. We made the tool as easy-to-use and as flexible as possible, and we don’t expect that the average user will require to add any functionalities. We think that one of possible limitations of the tool is that the researcher willing to add new functionalities to RTP2 will need to further expand her/his knowledge on Docker containers, server configurations and bash scripting.

There are important considerations for the researcher interested in using this tool or diffusion in general. For example, our results indicate that the WBT procedure, based on a whole-brain tractogram and white-matter ROIs, provided the most reproducible results. It is worth noting that these results come at a cost of precision and specificity. With the WBT procedure, there is limited control over the cortical endings of the tract, and it usually only works well with well-known big fascicles. For other smaller or more complicated tracts (e.g., dentatothalamic tract, optic radiation), the subcortical-cortical or cortico-cortical approaches provide slightly better results. Depending on the tract of interest, it is always worth testing in a couple of subjects to find out the best results. Additionally, we observed that the smaller the ROI is, the slightly worse are both the computational and test-retest reproducibility. This is not specific to our tool; as above, the researcher will need to decide how to combine the best ROIs and parameters to obtain the best possible tracts in the corresponding dataset/population. We plan to add as many recommendations as possible in the RTP2 wiki, so that researchers can conduct their own analyses at an advanced starting point (github.com/garikoitz/RTP-pipeline/wiki/Parameter-recommendations).

In conclusion, the RTP2 solution implements fully reproducible analyses, in an automated manner, while maintaining high levels of flexibility. In the Methods section we describe the computational steps involved, and some of the multiple possible configurations allowed. The methods we used guarantee that each tract profile can be traced to the original DICOMs. RTP2 implements reproducible computational methods by embedding algorithms on containers, which can be installed in a dedicated one-stop machine installed in the lab, or they can be installed in a HCP cluster, or other platforms such as Flywheel. In all cases, RTP2 will provide the same results if the input data is the same. Most importantly, this will be the case too if the data is reanalyzed now or in several years. We believe that RTP2 can be a helpful neuroimaging software tool consistent with modern requirements by being reproducible, automated/easy-to-use, and flexible.

## Acknowledgments

G. L-U. was supported by project IJC2020-042887-I funded by the Spanish Ministry of Science and Innovation; M.L. was supported by grants from the European Union’s Horizon 2020 research and innovation programme under the Marie Sklodowska-Curie (grant agreement No. 713673), and from “la Caixa” Foundation (grant No. 11660016); P.M.P.-A. was supported by grants from the Ministerio de Ciencia e Innovación (PGC2018-093408-B-I00), Neuroscience projects from the Fundación Tatiana Pérez de Guzmán el Bueno, Basque Government (PIBA-2021-1-0003), and a grant from “la Caixa” Banking Foundation under the project code LCF/PR/HR19/52160002. BCBL acknowledges support by the Basque Government through the BERC 2022-2025 program and by the Spanish State Research Agency through BCBL Severo Ochoa excellence accreditation CEX2020-001010-S.

We would like to acknowledge and express our gratitude to Leonardo Tozzi, Silei Zhu, Hongjian He and Lisa Bruckert for their help testing the tool, being patient with the bugs, proposing new functionalities, and in general for using it and making it better.

## Author contributions

Conceptualization, G.L-U. and B.W.; Methodology, Validation and Funding Acquisition, G.L-U., L.M., P.M.P-A., and B.W.; Resources, P.M.P-A., and B.W.; Software, G.L-U., L.M. and B.W.; Formal Analysis, Investigation and Visualization, G.L-U., and L.M.; Data Curation, G.L-U., L.M. and P.M.P-A.; Writing - Original Draft, G.L-U. and B.W.; Writing - Review and Editing, G.L-U., L.M., P.M.P-A., and B.W.; Supervision and Project Administration, G.L-U., P.M.P-A., and B.W.

## Declaration of interests

Brian Wandell works with Flywheel Exchange, LLC.

## Methods

Here we provide MRI data acquisition details (used in this paper as an example dataset) and describe the most important characteristics of RTP2. The small details, instructions about how to run it, and example configuration files are available in the wiki pages of the publicly available github sites (see below for links). We divided the description of the tool into three parts: (1) *the container*, (2) *the infrastructure* required to run them in a computationally reproducible system, the neuroinformatics platform (Marcus et al., 2011); and, (3) *the DWI data analysis pipeline*. The general idea is that regardless of the neuroinformatics platform of choice, the results will be the same because the computations inside of the containers will be the same.

### MRI data acquisition

A total of 112 healthy volunteers (mean age = 24.4 years, SD = 4.22 years; 64 females) participated in the study. Twenty-four of the volunteers (mean age = 24.7 years, SD = 4.06 years; 13 females) returned for a second session in which they were scanned using the exact same MRI protocol (mean interval = 15 days, SD = 21.82 days, range: 7-104 days). All participants were right-handed and had normal or corrected-to-normal vision. No participant had a history of major medical, neurological disorders, or treatment for psychiatric disorders. The study protocol was approved by the Ethics Committee of the Basque Center for Cognition, Brain and Language (BCBL) and was carried out in accordance with the Code of Ethics of the World Medical Association (Declaration of Helsinki) for experiments involving human participants. Prior to their inclusion in the study, all subjects provided informed written consent. Participants received monetary compensation for their participation.

Whole-brain MRI data acquisition was conducted on a 3-T Siemens Prisma Fit whole-body MRI scanner (Siemens Medical Solutions) using a 64-channel whole-head coil. The MRI acquisition included one T1-weighted structural image (T1w) and two DWI sequences. High-resolution MPRAGE T1-weighted structural images were collected with the following parameters: time-to-repetition (TR) = 2530 ms, time-to-echo (TE) = 2.36 ms; flip angle (FA) = 7°, field of view (FoV) = 256 mm, voxel resolution = 1 mm3, 176 slices. Both DWIs shared the following parameters: TR = 3600 ms, TE = 73 ms, FA = 78°, voxel resolution = 2 mm3, 72 slices with no gap and a multiband acceleration factor of 3. The first DWI acquisition of 105 volumes and anterior to posterior phase-encoding consisted of 5 blocks of 21 volumes: one without diffusion weighting (b0; b-value of 0s/mm2), 10 with a b-value of 1000 s/mm2 and 10 with a b-value of 2000 s/mm2, resulting in 50 isotropically distributed diffusion-encoding gradient directions for both the b=1000 and b=2000 shells. The five b0 images were acquired for motion correction. The second DWI acquisition of only seven b0 volumes was collected with reversed (posterior to anterior) phase-encoding to be used in geometrical distortion correction.

### Container creation and configuration

The container creation process starts in a client machine (e.g., laptop, personal computer or development server). After testing that our analysis tool works locally (anatROIs, RTP-preproc or RTP-pipeline in our case), we start the Docker Image creation process. Docker is the platform where the Docker images run, and a running Image is called a Container, i.e. a Docker Container is the run-time instance of a Docker Image. The first step is selecting a base Image that includes the required features. For example, as RTP-pipeline is a Matlab based program, we select a base Docker Image with the Matlab r2020b runtime, which is itself based on Ubuntu. In this base Docker Image, we install all the required programs, for example mrTrix (for an updated complete list of installed programs, refer to the Dockerfile file in https://github.com/garikoitz/RTP-pipeline). At this point, if it is a Matlab based program we compile it, otherwise, we build the Docker Image. Building the Image means obtaining all the required files and programs, including the compiled Matlab program if required.

The full cycle of testing a container has the following steps (in this example we show RTP2-pipeline, which uses Matlab, anatROIs and RTP-preproc are simpler as they do not use Matlab): (1) Run the code directly in Matlab and check the outputs; (2) Compile the Matlab code and run it using Matlab’s Runtime environment (it is freely downloadable from www.mathworks.com); (3) Build the Docker Container and run it using the default parameters and changed parameters; (4) Check the code in github, tag it, and build the containers in dockerhub; (5) Download the containers in another machine and check with test data that they run appropriately. At this moment, we can tag the container and upload to Dockerhub, and it will be available for anybody to use.

### Infrastructure for computational reproducibility

RTP2 is organized in 3 containers that can be run in a number of environments. They can be run natively on any computer if the required software is installed as well. We are going to illustrate three possible ways of running the pipeline (see Figure 1, bottom) using example cases.

#### Single computer

Examples of this computing platform are: (1) a single computer in the scanning room that we want to use for diffusion tractography, (2) a personal computer/laptop, (3) a small server in the lab/office we use for computing. In this case, we recommend having a Docker client implementation. Docker can be freely installed on a computer from https://www.docker.com/ by any user having administrator privileges. We tested it in macOS and Linux, and it should work in Windows too. The installation is typically simple, and it is done through the Docker client searching for the containers nipy/heudiconv, garikoitz/anatrois, garikoitz/rtppreproc, garikoitz/rtp-pipeline or using the command line:

~~~
docker pull nipy/heudiconv
docker pull garikoitz/anatrois
docker pull garikoitz/rtppreproc
docker pull garikoitz/rtp-pipeline
~~~

We recommend checking the latest version before installing. Once the containers are installed they are ready to run. For specific instructions about how to launch the containers, please see the wiki.

For individual subjects, it is manageable to launch the containers manually in the command line. This would be the recommendation for the testing phase. In the operational phase, we recommend a data organization and container launching scheme programmed in python. It can be obtained here https://github.com/garikoitz/launchcontainers. The instructions on how to use it are in the wiki of the repository. In order to save space, it creates links to files and takes care of the versioning of the files. Once the containers end the computations, the same computer can be used for tract visualization, quality assurance, or statistical analyses.

#### High Performance Computing (HPC) Systems

It is common for many research labs to have access to HPC systems. These systems will be maintained by IT professionals, and typically will provide the researchers/clinicians with a way of accessing the system and running their data. We have not encountered any of those systems that allow running Docker containers, but all of them accept running Singularity containers. The procedure is almost the same as for the Docker containers. Once the DICOMs files from the scanner have been transferred to the HPC system, it is possible to run our results there and copy the results back to our visualization and statistical analysis computers.

Installation is as simple as with the Docker containers. Importantly, it does not require assistance from IT professionals. These are the commands that would install the containers in the home folder of the researcher:

~~~
singularity build $ HOME/heudiconv.sif docker://nipy/heudiconv
singularity build $ HOME/anatrois.sif docker://garikoitz/anatrois
singularity build $ HOME/rtppreproc.sif docker://garikoitz/rtppreproc
singularity build $ HOME/rtp-pipeline.sif docker://garikoitz/rtp-pipeline
~~~

Once installed, we recommend using the same python code in https://github.com/garikoitz/launchcontainers to prepare the data and launch the containers, although it is possible to launch everything manually. Please refer to the wiki for more instructions.

#### Cloud based neuroinformatics platform (Flywheel.io)

Cloud based systems and software-as-a-service (SaaS) systems are taking over the software service landscape. The analysis containers were developed (and are extensively used) to be used as part of the neuroinformatics platform called Flywheel. If your institution provides a subscription, you will be able to install these containers and use them as part of an integrated cloud based tool. Flywheel.io implements reproducible computational methods, tracks provenance of the data, and facilitates data sharing. The main functionalities provided by Flywheel are: *(1) Data Capture:* automatic data conversion and uploading from the scanner, *(2) Data Curation:* visualization, computation, sharing, download, reorganization, sharing, and renaming, can be done with remote tools (for example https://github.com/vistalab/scitran/), (3) *Compute:* running containers; every analysis, including versions, parameters, inputs, and outputs, is named, time-stamped and stored in the searchable database. (4) *Collaborating:* all the data, metadata and analyses can be made available to anyone provided with permission in the organization. Containers are called gears in Flywheel. The same containers created in the first section are used as gears. For a detailed Gear creation and installation process, please refer to Flywheel’s official documentation in https://docs.flywheel.io. See Figure S4 for details in Flywheel’s technical architecture.

An important component of the whole RTP2 solution is that it allows programmatic access and control of the whole system. The same thing is true even if we run RTP2 in the Flywheel system. We provide Matlab and Python based software similar to the *launchcontainers* software to control the cloud engine. In Flywheel the data storage and the computation jobs are always done in the cloud server. For a few subjects the interaction with Flywheel can be done through the web interface and most of the settings and tools can be tested this way. But increasingly, neuroimaging projects consist of a larger number of subjects, and programmatic access is required for time efficient and error free data, configuration parameters and result management. Even with two subjects, it is easy to make mistakes when setting the more than 50 RTP2 config parameters. To solve this problem we developed the Scitran tool. Scitran is a Matlab application built on top of Flywheel’s Matlab SDK, that provides several functionalities: *(1) Data/analysis results search:* to perform any of the following operations, *(2) Metadata update:* for example, socioeconomic information from the subjects, *(3) Upload and download information:* log files, images, or other project level required information, *(4) Job launch and management:* select a gear, version, config parameters and launch an analysis job and control the status, *(5) Information extraction:* select the desired subjects and analysis and extract only the required results that go to the statistical analysis. Specifically for RTP2, there is a utility that extracts the per-subject-per-tract metric (e.g. FA) and creates a table with all the project level, acquisition level and analysis level parameters. Scitran is publicly available and can be installed from (https://github.com/vistalab/scitran/). Installing it in Matlab is as simple as downloading the zip file or cloning the repository and adding it to the Matlab path. Scitran comes with the latest Flywheel Matlab SDK. In order to make it work, it is required to have an active user in a Flywheel instance and download an API key. Please refer to the Scitran manual (https://github.com/vistalab/scitran/wiki) for further information on installing and using the Scitran tool.

### Data analysis pipeline

#### anatROIs

The first step of the RTP2-pipeline (called RTP2-anatROIs) involves the processing of the subject’s anatomical T1w image, with the objective of obtaining the ROIs that will later be used in tractography. The input of this step is the subject’s T1w file and ROIs defined in MNI space; the output is a segmented T1w image and our ROIs of interest in individual subject T1w space. AnatROIs provides many predefined ROIs, and it is possible to extend it in multiple ways (see Figure 2; https://github.com/garikoitz/anatROIs/wiki).

##### Included ROIs

the T1w image is processed with the Freesurfer package (http://surfer.nmr.mgh.harvard.edu/) to obtain a cortical/subcortical segmentation and parcellation. If a previous freesurfer run exists, it is possible to use it, just passing the folder or a zip file with the completed analysis. This step already provides all of Freesurfer’s segmentations and parcellations (check freesurfer or the wiki for more detail). Importantly, the surface parcellations are converted automatically to volumetric ROIs so that they can be used in tracking automatically. To convert ROIs or atlases defined in MNI space to individual-subject space, the container performed a non-linear registration of a 1mm^3^ MNI template using Advanced Normalization Tools (ANTs, http://stnava.github.io/ANTs/). On top of all the Freesurfer segmentation (aseg, hippocampus, thalamus, brainstem) and parcellation (aparc2009) tools, we provided several ROIs in MNI space, the human connectome project (HCP) atlas (Glasser et al., 2016) and cerebellum parcellation proposed by (Diedrichsen et al., 2011). To parcellate the visual cortex we run the Neuropythy (Benson and Winawer, 2018) tool on top of Freesurfer’s results. Check the wiki for the latest version’s available ROIs and segmentation tools (https://github.com/garikoitz/anatROIs/wiki/How-to-use).

##### User generated ROIs

there are many ways to add ROIs or atlases to anatROIs. *(1) In template space:* MNI ROIs can be directly added to the container, as well as surface ROIs drawn in Freesurfer’s fsaverage. *Native space:* similarly, ROIs can be drawn in native space, either in 3D space or in the surface. The ROIs can come from analyses in other modalities, for example fMRI results. Please refer to the wiki for more details (https://github.com/garikoitz/anatROIs/wiki).

#### RTP2-preproc

The preprocessing container prepares the data for the posterior DWI analysis in the RTP2-pipeline. The preprocessing does not include data analysis: its objective is to correct known problems in the DWI acquisitions process. It can optionally align the DWI data with the anatomical file (done by default). This gear is an adaptation from mrTrix’s (https://github.com/MRtrix3/mrtrix3) recommended preprocessing procedure implemented in Brainlife (https://brainlife.io/app/5a813e52dc4031003b8b36f9). This container takes some required and optional inputs. More details in the wiki of the repository: https://github.com/garikoitz/RTP-preproc/wiki.

- Required:
  - Diffusion data in 4D Nifti format
  - Bvec file in FSL format
  - Bval file in FSL format
- Optional:
  - Anatomical T1w file: used to optionally align the diffusion data to it
  - Reversed phase encoding DWI data.
  - Bval file, corresponding to the reversed phase encoding DWI data.
  - Bvec file, corresponding to the reversed phase encoding DWI data.
  - Brainmask: it can be Freesurfer’s brainmask or any mask edited manually. If a mask is provided, the container does not use FSL’s BET tool, which usually doesn’t give very good results.

It is highly recommended to acquire the reversed phase encoding files to correct for EPI geometric distortions with FSL’s TOPUP. If those images are not provided, the alignment between the diffusion data and the anatomy is not going to be as good. This can be relevant downstream for analysis requiring a good alignment between anatomical and diffusion data —for example if we want to use mrTrix’s Anatomically Constrained Tractography, or any cortical ROI for that matter.

In the wiki we provide an exhaustive list of the preprocessing steps that can be set using the configuration parameters (https://github.com/garikoitz/RTP-preproc/wiki/configjson). We will use an application logic order here, this is the chronological order of how the options are applied. Once the analysis is run, it is recommended to visually inspect the results, if not in all the subjects, in a representative number of those, to avoid any systematic and database dependent errors.

#### RTP2-pipeline

The tractography container takes preprocessed DWI data from RTP2-preproc and the ROIs from anatROIs and automatically identifies any number of user-specified white matter tracts. Using the ROIs and atlases provided by default (check the wiki for an updated list https://github.com/garikoitz/anatROIs/wiki/atlases) or by adding new used defined ROIs, the tracking possibilities are endless. Once the tracts are identified, it provides the tract and its metrics in several different ways (see Figure 5). In the analysis process, the pipeline generates values at the voxel and the tract level, which can be later used for further analysis.

**Figure 5.**
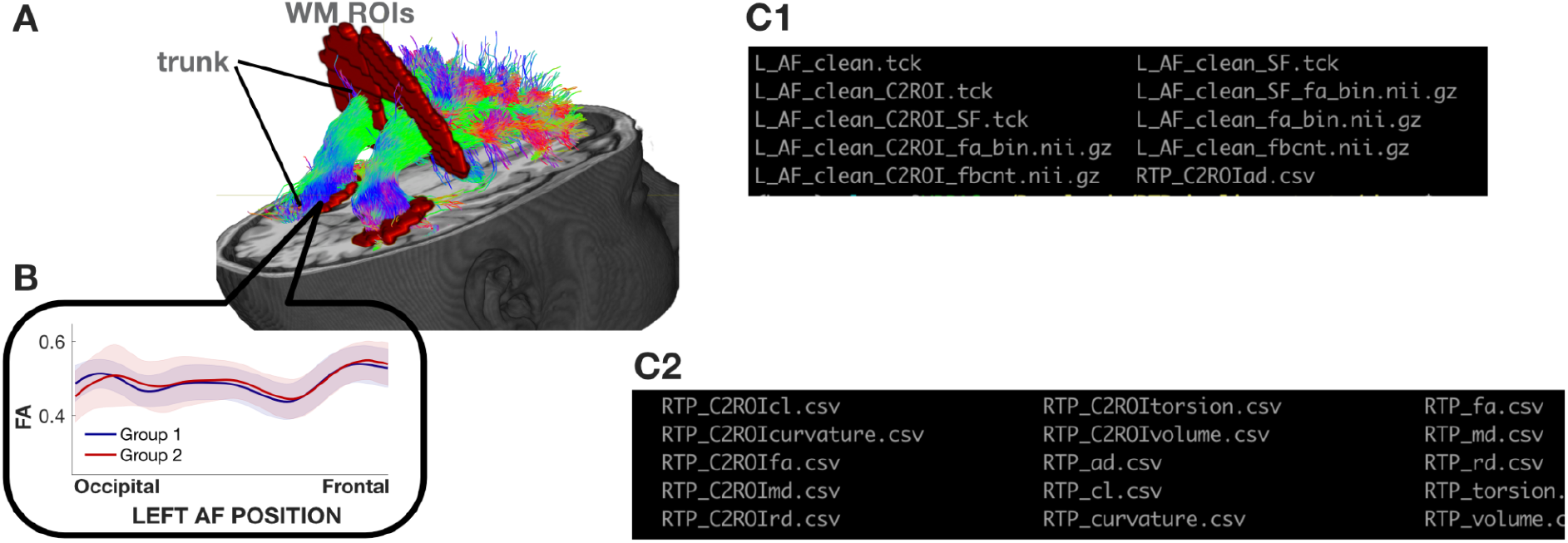
Illustrative tract (arcuate fasciculus) with defining ROIs and tract profile. **A**. The streamlines serve as a model of white matter tracts; they are selected by fitting to the diffusion-weighted imaging (DWI) measurements. The tracts are defined by regions of interest (ROIs, red) that select specific streamlines from the whole brain tractogram (in this example). The region between the two ROIs is relatively stable and called the trunk. **B**. Tract’s FA profile. We can calculate profiles from the whole tract or only the trunk, called clip-to-roi (C2ROI) operation. We estimate a core fiber (called super fiber —SF) from the collection of streamlines and sample 100 equally spaced segments. The FA of the core fiber is calculated by combining FA transverse to the core fiber at every sample point, using a Gaussian weighting scheme over distance. The set of sample points is the tract profile; the average of the FA values of the core fiber is the mean tract FA. In the inset, an illustrative example of two group profiles. The dark outline is the mean of the group and the shaded outline represents a standard deviation of the values. **C1**. List of files in the output of RTP2-pipeline. Every tract generates a number of different outputs, for both the C2ROI and normal options. Tck files are tract definitions, and they can be the full tract or only the super-fiber (marked as _SF). On top of the tck-s, we created two types of nifty, the fa_bin Nifti is a mask of the tract, with 1-s where there are fibers and 0-s elsewhere. In the fbcnt Nifti it is zero where there are no fibers, and in the rest of the voxels the value is the fiber count in the voxel. **C2**. The csv files contain the tract profiles for all of the different tracts that were tracked in this specific subject. There is one file per metric. Thus, the size of every csv is M^row^ x N^column^ (M: 100, N: number of tracts obtained in this analysis).

There are two main approaches to generate one tract: edit from whole brain streamlines, or generate originally from given ROIs. The recommendation is to apply the first approach when possible, because whole-brain streamlines can be validated and trimmed by specific algorithms (see details described next). In the current version (RTP2), RTP2-pipeline is heavily based on mrTrix. The list of parameters is too long to show it here, for a complete list please refer to the manifest.json file in (github.com/garikoitz/RTP-pipeline/wiki). The wiki contains an example_config.json file with all the default values (for Flywheel, the manifest.json contains a description and a default value per every parameter).

It is important to notice that the parameter selection is project specific. We recommend using the default in a first test run, but from that moment on, it is usually required to adjust the parameters. The github repository contains a wiki page with parameter recommendations (github.com/garikoitz/RTP-pipeline/wiki/Parameter-recommendations) for different types of acquisition data, with the hope that it will be of help to other researchers. We hope that this list of recommendations will grow in the future.

This container has a list of required files:

- Preprocessed diffusion data (output from RTP2-preproc, ideally)
- Anatomical T1w image (output from anatROIs, ideally)
- Zip file containing the ROIs (output from anatROIs, ideally, but it can be created manually, or use any)
- Config.json file: it will be used mostly for the whole tractography
- Tractparams.csv file: it indicates all of the tracts that we want to obtain, detailing the ROIs and the configuration parameters. See github.com/garikoitz/RTP-pipeline/wiki/How-to-edit-tractparams.csv for an example and instructions on how to fill it. If a new acquisitions sequence or special population is being used, it will require adjustments the first time it is run. The ROI names used here refer to the ROI names in the zip file.

Next, we illustrate the main steps implemented in the pipeline and will explain the main configuration parameter choices:

1. ROI validation. Check existence of ROI files, and dilate/combine ROIs if specified in the tractparams.csv file (see github.com/garikoitz/RTP-pipeline/wiki/How-to-edit-tractparams.csv).
2. DTI modeling at the voxel level and metric creation: FA, MD, RD etc.
3. Response function creation. Different algorithms can be selected, and with different options. We recommend using the dhollander algorithm with automatic lmax, which separates the WM, GM, and CSF in the DWI without requiring an anatomical file.
4. Constrained Spherical Deconvolution (CSD) modeling. Different algorithms can be selected. Performed at the voxel level, it creates Fiber Orientation Distributions (FOD) that are required for tracking.
5. Whole-brain white-matter streamlines estimation. Whole-brain streamlines will be estimated when there is at least one tract being generated from editing whole-brain streamlines (as specified in tractparams.csv file). Different algorithms and options are available (see mrTrix documentation for a comprehensive list on the tckgen command). Additionally, there are other decisions that can be taken at this point. We implemented Ensemble Tractography (Takemura et al., 2016), SIFT (Smith et al., 2013) and LiFE (Linear Fascicle Evaluation) (Pestilli et al., 2014) into the RTP2-pipeline. It is possible to run them independently, i.e. both, one or the other, or none can be selected.
  a. Ensemble Tractography (ET) method: ET invokes MRtrix’s tractography tool a number of times (selected with a configuration parameter), constructing whole-brain tractograms with a range of tracking parameter options. For example, the minimum angle parameters for tracking can be varied (e.g. 47, 23, 11, 6, 3), and the resulting whole-brain connectomes concatenated together. There are config parameters specific to the Ensemble Tractography implementation.
  b. The SIFT and LiFE (Linear Fascicle Evaluation) methods evaluate the tractogram streamlines and retain those that meaningfully contribute to predicting variance in the DWI data. There are config parameters specific to the SIFT and LiFE implementation.
6. Tracts generation. User-defined tracts can be defined either from selecting specific streamlines from the whole-brain tractogram (see previous step), or generating streamlines using tckgen in MRtrix using seed and target ROIs.
7. Tract Profiles. The voxel-based metrics are calculated for each individual tract creating the tract profiles. The metrics can be of any voxel-based measurement. Although the most common ones are the DTI metrics (FA, MD, …), we have successfully used others, such as T1 relaxation time or macromolecular tissue volume (MTV) quantitative MRI maps. The basic steps for tract profile creation are:
  a. A core fiber, representing the central tendency of all the streamlines in the tract, is identified.
  b. Equally spaced positions along the fiber between the two defining ROIs are sampled (N=100).
  c. The metric values of streamlines at locations transverse to each sample position are measured and combined. The combination is weighted by a Gaussian value based on the distance from the sample point.
  d. The sampling and transverse averaging generates a tract profile of 100 metric values (see inset in Figure 4 for a group mean FA tract profile).
8. Tract cleaning. After generating the tract, and calculating the core fiber, it is possible to clean the tract by removing any streamlines that (1) have length that are specific standard deviations away from the mean fiber length or (2) the distance to the core fiber fell out of specific standard deviations from the mean distance of all streamlines. This means that fiber groups will be forced to be a compact bundle.
9. Export tracts as Nifti masks (see <u>Figure 6-C1</u>). It creates volumetric images in native and template space, creating binary masks (1 if a fiber crosses the voxel, 0 if not), or fiber count masks (each voxel indicates the amount of fibers crossing the voxel).

## Supplemental information

**Figure S1.**
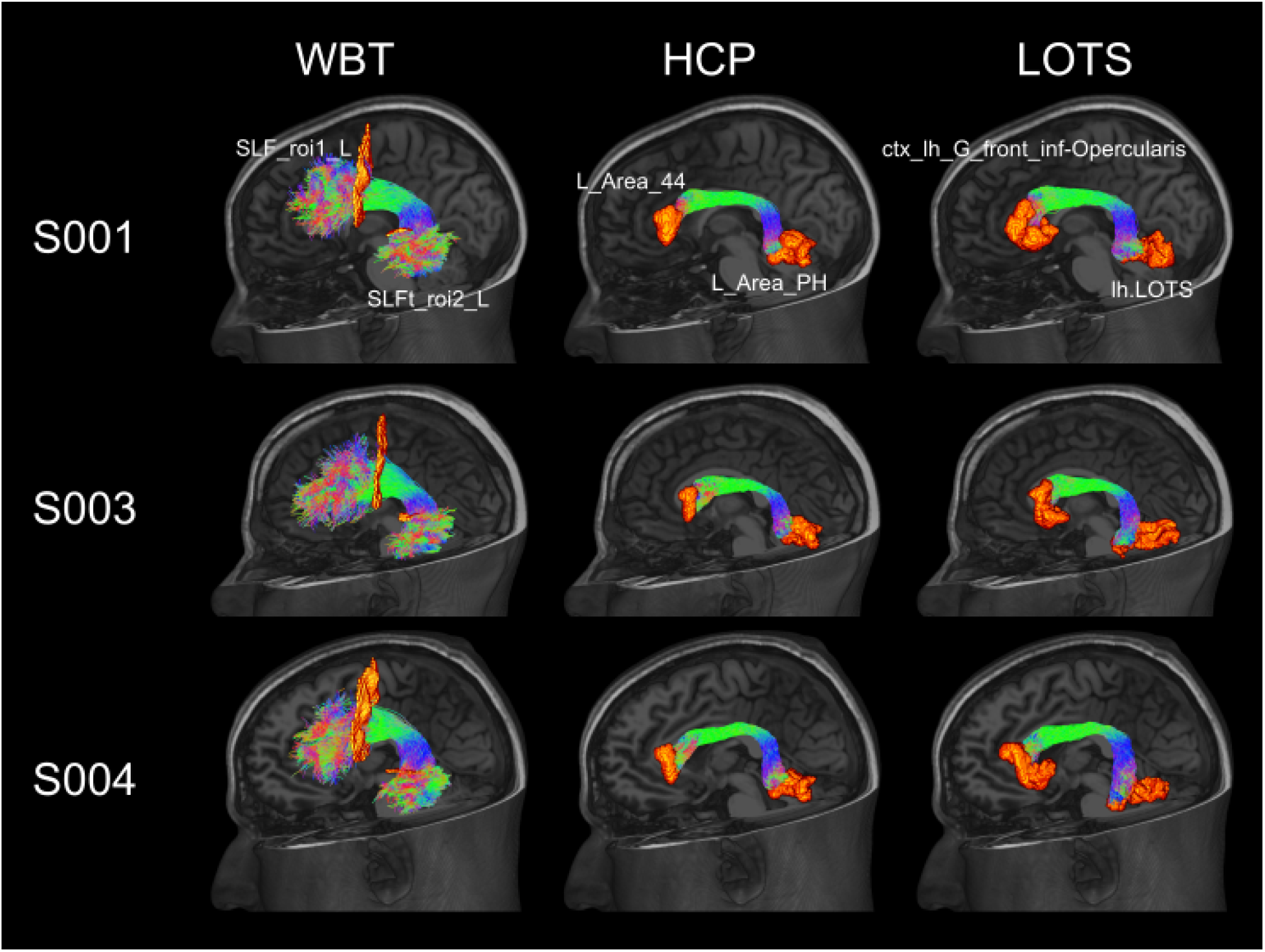
Rendered image of left arcuate fasciculus obtained with three different procedures in three representative brains. In the WBT procedure, the left AF is obtained by selecting streamlines from the whole-brain-tractogram passing through white matter ROIs. In the HCP procedure, the left AF is obtained by directly tracing between area 44 and area PH from the HCP atlas. In the LOTS procedure, similar to the HCP procedure, the left AF is obtained by directly tracing between two cortical ROIs, opercularis obtained from Freesurfer’s cortical atlas, and LOTS manually drawn in Freesurfer’s fsaverage surface template. All ROIs are represented in orange, and all fibers are represented in colors attending their direction.

In addition to FA, we measured the reproducibility of tract profiles using other microstructural tractometric indices: axial diffusivity (AD), mean diffusivity (MD) and radial diffusivity (RD). Similar to the reproducibility obtained for FA values, all three indices showed high computational (Supplementary Figure S2) and test-retest reproducibility (Supplementary Figure S3) across all tracts of interest. In test-retest reproducibility, as expected, the correlation coefficients tended to be numerically lower relative to computational reproducibility, but they still show high reproducibility at the group level.

**Figure S2.**
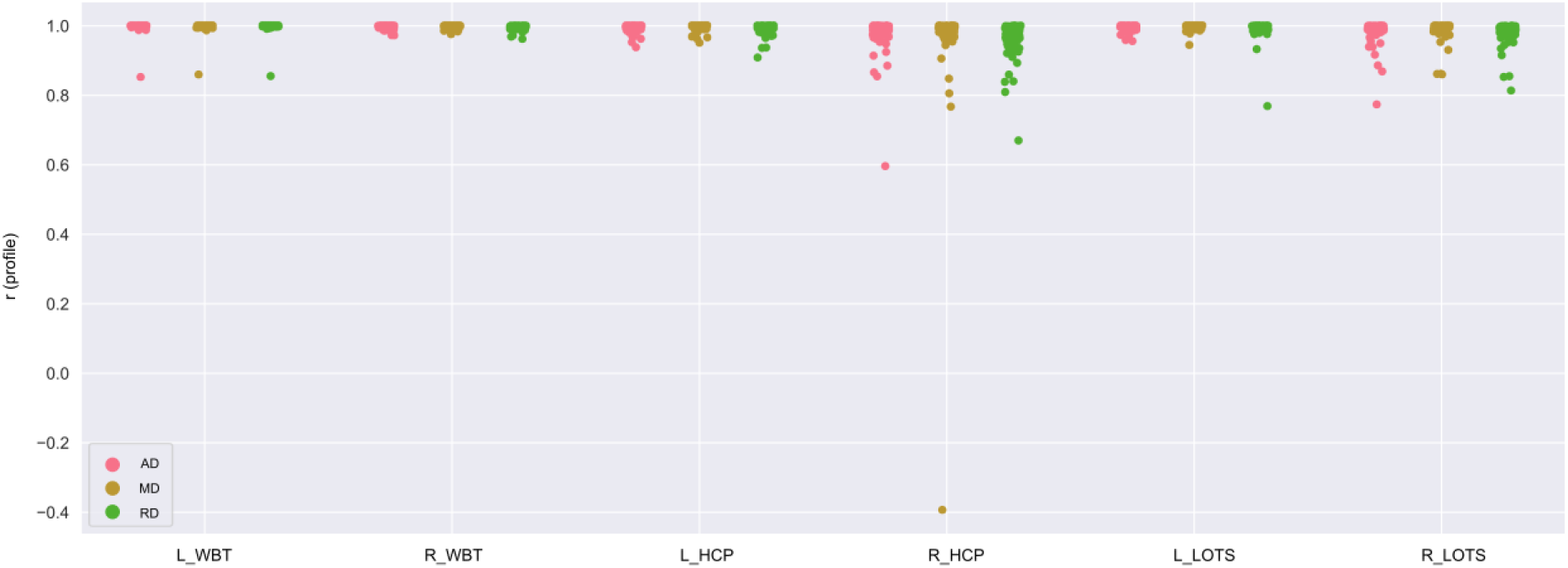
Computational reproducibility of fiber profiles for several microstructural measurements. Correlation coefficients of fiber profiles on axial diffusivity (AD), mean diffusivity (MD) and radial diffusivity (RD) between two separate computations. Each dot represents one instance of the correlation coefficient of one participant between computations.

**Figure S3.**
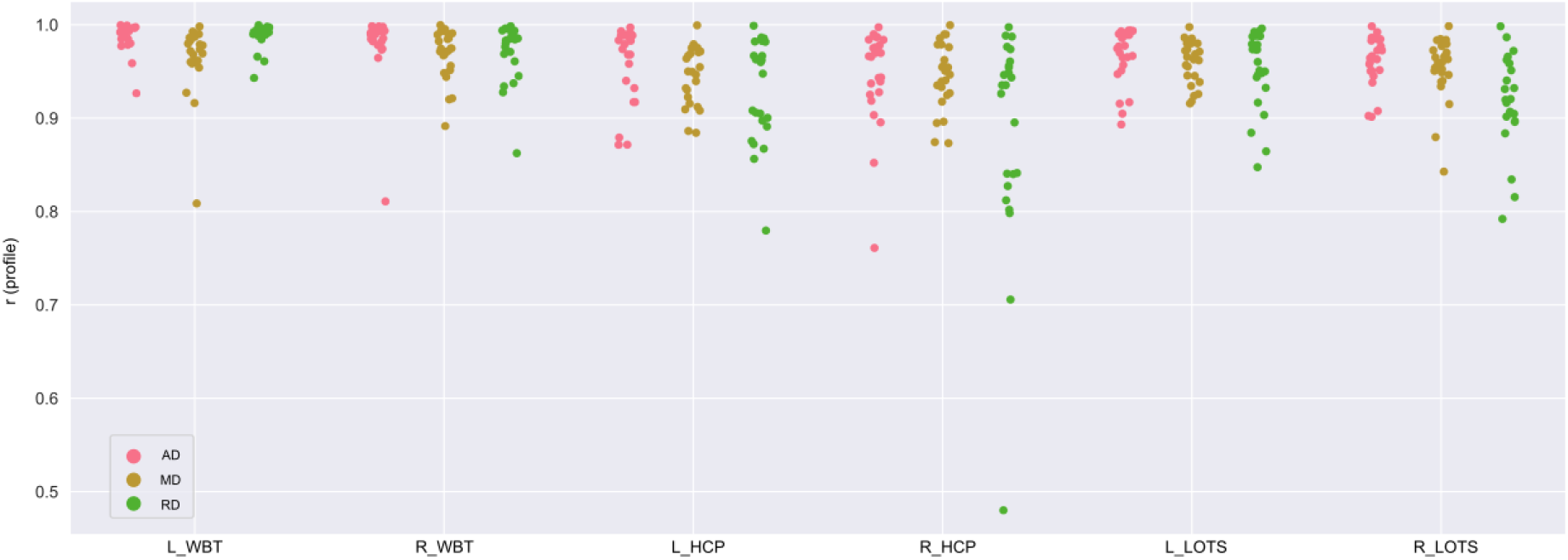
Test-retest reproducibility of fiber profiles for microstructural measurements. Correlation coefficients of fiber profiles on AD, MD and RD between test and retest computations. Each dot represents one instance of the correlation coefficient of one participant between test and retest computations.

**Figure S4.**
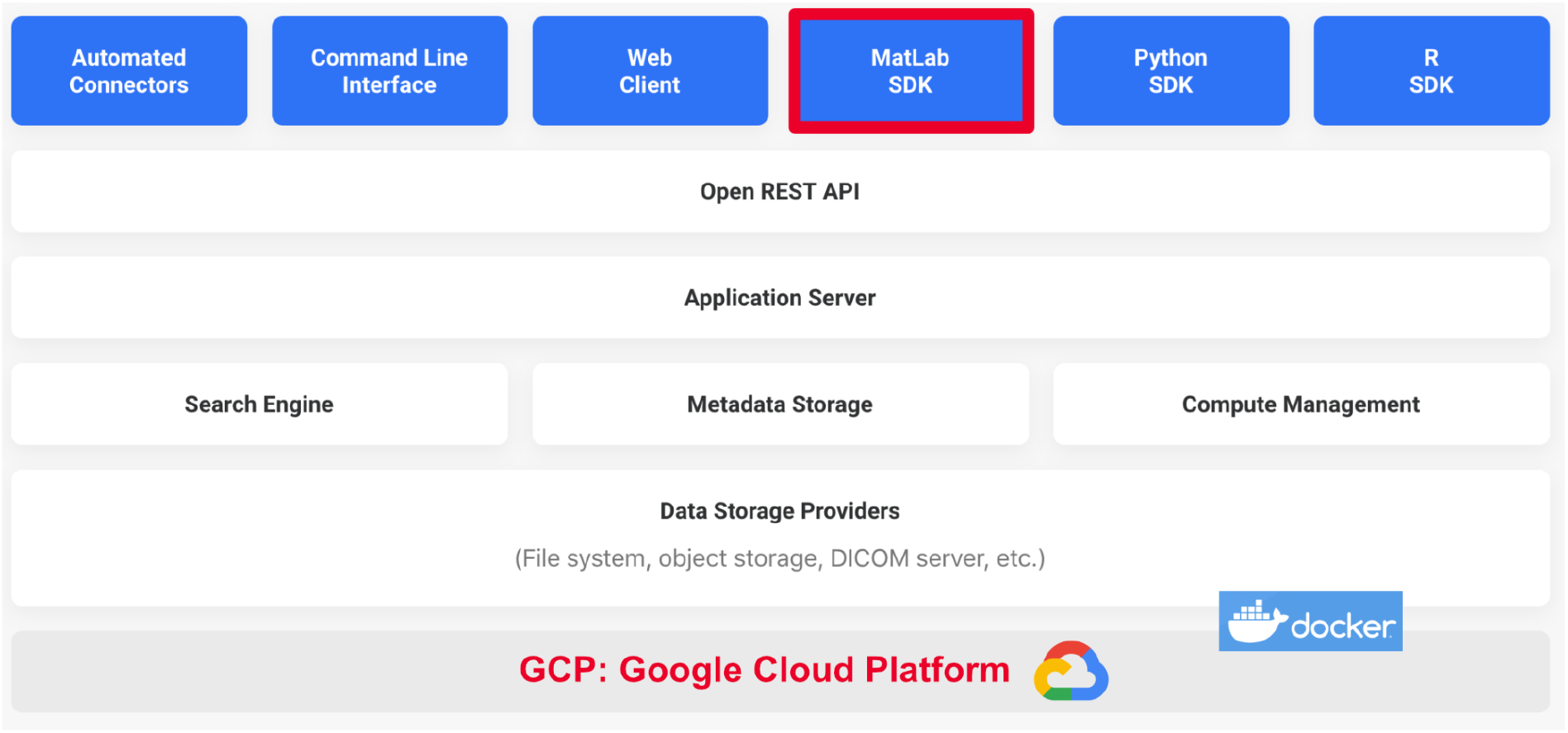
Flywheel’s technical architecture. The first row (blue) shows the different ways to interact with the system: *(1) Automated connectors:* load data from the MRI scanners to Flywheel. *(2) Command Line Interface:* a program installed locally allows the authentication to the Flywheel system and most of the management operations. Useful, for example, to upload existing BIDS formated datasets to Flywheel if we don’t have access to the original scanner. *(3) Web Client:* web interface to the Flywheel system accessible from all the major browsers. Allows the majority of the operations with a clean point and click interface. *(4-5-6) Matlab-Python-R SDK:* the SDK gives programmatic access to the Flywheel system, with the three main scientific programming languages. We marked Matlab SDK in red in Figure 5 because as part of the RTP2 solution we included Scitran (https://github.com/vistalab/scitran), which is a collection of Matlab scripts designed to interact programmatically with Flywheel. These interfaces use the Open (public) REST API (Representational State Transfer Application Programming Interface) to access Flywheel’s Application Server, which control the web interface and Flywheel’s three main systems: *(1) search engine:* accessible through the web GUI and from the SDK to search and retrieve only the required information, *(2) Metadata Storage:* core of the system, with references to all content, gear and analyses, and *(3) Compute management:* it manages the dispatching of jobs, i.e. it takes the input data, runs the Gear, and when finished, copies the results back to Flywheel. The last two rows represent the data storage and the actual hardware where Flywheel is running (and where the Gears run as well). In our specific instance of Flywheel, both the data and the computations happen in the Google Cloud Platform. Small Gears run on the same hardware as Flywheel, but big Gears (as all three gear composing RTP2) launch a new virtual machine each time, runs the Gear, and shuts it off when finished. There is a parameter in Flywheel that controls the number of machines that can run in parallel. This is required for both technical and economical reasons, as Google is not usually the bottleneck in this situation.

